# A hidden battle in the dirt: soil amoebae interactions with *Paracoccidioides* spp

**DOI:** 10.1101/564435

**Authors:** Patrícia Albuquerque, André Moraes Nicola, Diogo Almeida Gomes Magnabosco, Lorena da Silveira Derengowski, Luana Soares Crisóstomo, Luciano Costa Gomes Xavier, Stefânia de Oliveira Frazão, Fernanda Guilhelmelli, Marco Antônio de Oliveira, Fabián Andrés Hurtado, Marcus de Melo Teixeira, Allan J. Guimaraes, Hugo Costa Paes, Eduardo Bagagli, Maria Sueli Soares Felipe, Arturo Casadevall, Ildinete Silva-Pereira

## Abstract

*Paracoccidioides* spp. are thermodimorphic pathogenic fungi endemic to Latin America. Predation is believed to drive the evolution of virulence for soil saprophytes. We evaluated the presence of environmental amoeboid predators in soil from armadillo burrows where *Paracoccidioides* had been previously detected and tested if interaction of *Paracoccidioides* with amoebae increased fungal virulence. Nematodes, ciliates and amoebae – all potential predators of fungi – grew in cultures from soil samples. Microscopical observation and ITS sequencing identified the amoebae as *Acanthamoeba* spp, *Allovahlkampfia spelaea* and *Vermamoeba vermiformis*. These three amoebae efficiently ingested, killed and digested *Paracoccidioides* spp. yeast cells, as did laboratory-adapted axenic *Acanthamoeba castellanii*. Sequential co-cultivation of *Paracoccidioides* with *A. castellanii* selected for phenotypical traits related to survival of the fungus within a natural predator as well as in murine macrophages and in vivo (*Galleria mellonella* and mice). This increase in virulence is linked to the accumulation of cell wall alpha-glucans, polysaccharides that masks recognition of fungal molecular patterns by host pattern recognition receptors. Altogether, our results indicate that *Paracoccidioides* inhabits a complex environment with multiple amoeboid predators that can exert selective pressure to guide the evolution of virulence traits.

## Introduction

Human beings are constantly challenged by microorganisms in virtually every environment and circumstance. Effective host immune responses, however, ensure that very few of them cause disease. Pathogenic microorganisms usually have a complex set of virulence attributes that allow them to evade immune effectors, proliferate and cause diseases [1]. Immunity is a crucial selective pressure driving the evolution of virulence attributes in microbial pathogens tightly associated with mammalian hosts. However, the evolution of virulence in microbes that do not need to interact with mammals to complete their life cycles, such as the agents of most fungal invasive diseases, is far less clear. These agents include pathogenic species in the genus *Paracoccidioides*. Five species in this genus of thermodimorphic fungi, *P. brasiliensis, P. americana, P. restrepiensis, P. venezuelensis* and *P. lutzii*, cause paracoccidioidomycosis (PCM), one of the most prevalent systemic mycoses in Latin America [2, 3]. This neglected disease is an important cause of morbidity and mortality among men from rural areas in these countries. Infection occurs by the inhalation of airborne fungal propagules (mycelium fragments or conidia) from the environment, and most infections are asymptomatic. However, some patients do develop disease, which ranges from a mild pneumonia to life-threatening systemic disease [4, 5].

In the last two decades, a number of studies done with other species of invasive fungal pathogens (*Cryptococcus neoformans, C. gattii, Sporothrix schenckii, Blastomyces dermatitidis, Histoplasma capsulatum* and *Aspergillus fumigatus*) have provided a compelling explanation for the evolution of virulence in these soil saprophytes: avoiding predation by soil amoebae requires phenotypical traits that also provide protection against mammalian immune defenses and are thus associated with virulence [6–10]. Each of these fungi survived after co-cultivation with phagocytic unicellular organisms such as *Acanthamoeba castellanii* and *Dictyostelium discoideum* due to phenotypical traits that are also effective in evading human macrophages. Moreover, their cocultivation with amoebae selects for survivors that are more virulent in mammalian models. Exposure to other soil predators such as ciliates and helminths suggests a more complex interaction scenario, beyond those seen with amoebae [11, 12]. These studies, however, were performed in controlled laboratory conditions using mostly pure and axenic cultures of laboratory-adapted predators; this very informative system is nonetheless an extreme simplification of the complex ecosystem soil saprophytes find in nature. In this work we have delved further into the ecology of the soil environment in a region where *Paracoccidioides* spp. had previously been confirmed by nested PCR, studying both the composition of predator populations and the interaction between some of these and *Paracoccidioides* cells. Cultures of the soil samples revealed helminths, ciliates and multiple species of amoebae. We successfully isolated amoeba species and showed that they can ingest and efficiently kill *Paracoccidioides* spp. yeast cells. The same was observed in more detailed experiments with axenic cultures of *A. castellanii*. Sequential co-cultivation of *Paracoccidioides* cells with *A. castellanii* selected for fungal cells with increased virulence towards both phagocytes (amoebae and ex vivo mouse macrophages) and whole animals (*Galleria mellonella* and mice), possibly due to an increase in cell wall alpha-glucans. Our results support the hypothesis that interaction with sympatric soil predators selects for traits that allow survival of *Paracoccidioides* spp. in mammalian hosts and add to the existing evidence for the amoeboid predator-animal virulence hypothesis [13].

## Materials and methods

### *Paracoccidioides* spp. strains

For our studies we used the *P. brasiliensis* clinical isolated isolate Pb18, *P. brasiliensis* isolate T16B1, isolated from the spleen of a nine-banded armadillo (*Dasypus novemcinctus*) [14] and the *P. lutzii* isolate Pb01. The yeast phase of these isolates was maintained and prepared for interaction with soil amoebae as described in Supplemental Materials and Methods.

### Axenic amoebae

*A. castellanii* 30234 (*American Type Culture Collection* - ATCC, Manassas, VA, USA) was cultivated in PYG medium (2% Proteose peptone, 0.1% yeast extract, 1.8% glucose, 0.1% Sodium citrate dihydrate, 2.5 mM Na_2_HPO_4_, 2.5 mM KH2PO4, 4 mM MgSO_4_, 400 μM CaCl_2_, 50 μM Fe(NH_4_)_2_(SO_4_)_2_) at 28°C as previously described [8].

### Soil amoeba isolation and maintenance

Soil amoebae were isolated from samples of armadillo burrows located at Lageado Farm (−22° 50’ 14.36″ latitude and −48° 25’ 31.35^y^ longitude), an area where an armadillo was captured and PbT16B1 was isolated; in this location the fungus had been also detected in soil by nested PCR [14]. Additionally, rural workers that have lived and/or worked in this region were diagnosed with or died from PCM [15]. About five grams of each soil sample were mixed with 20 mL of sterile Page’s modified Neff’s amoeba saline (PAS – 2 mM NaCl, 33 mM MgSO_4_, 27 mM CaCl_2_, 1 mM Na_2_HPO_4_, 1 mM KH_2_PO_4_) and vigorously mixed to homogenize the samples. After sedimentation 100 μL of each sample were spread over a plate of non-nutrient agar (PAS + 1.5% agar) containing a lawn of heat-killed *Escherichia coli* OP50. The plates were incubated at 25 °C for 10-14 days and observed daily by light microscopy for the presence of cysts or trophozoites of amoebae [16, 17]. Agar sections containing cyst or trophozoites were cut and transferred to new plates to enrich the cultures. Finally, amoebae were transferred to PAS, counted and submitted to limiting dilution cloning. These freshly isolated amoebae were maintained in PAS or in nonnutrient agar plates with live or dead *E. coli* strain OP50 as a food source, respectively. The isolates were molecular typed for identification as described in the Supplemental Materials and Methods section.

### Soil amoeba and *P. brasiliensis* interaction assays

The distinct amoeba isolates were collected from our culture stocks, washed three times to remove bacteria, plated onto glass-bottom plates and co-incubated with *P. brasiliensis* cells previously dyed with FITC or pHrodo™ (Thermo Fisher). The multiplicity of infection (MOI) was of one and co-incubation was carried out for 24 hours at 25 °C). The samples were then dyed with Uvitex to distinguish intracellular and extracellular fungal cells and observed in a Zeiss Axio Observer Z1 inverted microscope using a 40X/NA 0.6 objective for quantification of phagocytosis. A minimum of 100 amoebae per sample were analyzed, and the experiments were performed at least three times on different days. Alternatively, predation assays in which soil amoebae were incubated in solid non-nutrient agar with a lawn of *P. brasiliensis* cells were performed as described in the Supplemental Material and Methods section. Soil amoeba viability after the interaction was assessed by Trypan blue exclusion as previously described [8].

### Phagocytosis and killing assay for the interaction of *Paracoccidioides* spp with axenic *A. castellanii*

Cells of *A. castellanii* were plated onto 96- or 24-well microplates at 5 × 10^4^ and 2× 10^5^ cells/well, respectively, and incubated with yeast cells (1 × 10^5^ and 4 x 10^5^ cells/well) for different time intervals (6, 24 or 48 h) at 28 °C or 37 °C (MOI = 2). After co-incubation, the supernatant was discarded, the cells were fixed with cold methanol for 30 min at 4 °C and overnight-stained with Giemsa. The samples were then observed and photographed in the Zeiss Axio Observer Z1 inverted microscope. Alternatively, phagocytosis was evaluated using fungal cells previously dyed with CMFDA or FITC before the interaction. At each condition the percentage of phagocytosis was evaluated after Giemsa staining. A minimum of 100 amoebae per sample were analyzed, and the experiments were performed at least three times on different days. *A. castellanii* viability after the interaction was assessed by Trypan blue exclusion as previously described [8]. Fungal survival after the interaction was assessed by CFU counting after amoeba lysis as described in the Supplemental Materials and Methods section.

### Microscopical analysis of *Paracoccidioides* spp interaction with amoebae

We further analyzed the interaction between different soil amoeba with *Paracoccidioides* spp using confocal microscopy, Transmission Electron Microscopy (TEM) and Scanning Electronic Microscopy (SEM). The detailed approach for each technique is described in the Supplemental Materials and Methods section.

### Sequential passages of interaction of *A. castellanii* and *Paracoccidioides* spp

We co-cultured *Paracoccidioides* spp. cells with *A. castellanii* for six hours at 28 °C in PYG medium at a MOI of two. The cells were then detached from the plates and passed 5-8 times through a 26-Gauge syringe to lyse amoebae. The remaining yeast cells were plated onto solid BHI-Sup (4% horse serum, 5% conditioned medium of the Pb192 strain of *P. brasiliensis*, 34 μg/mL chloramphenicol). The plates were incubated at 37 °C for a week, and the recovered cells were collected from the plates, washed three times with PBS, counted and used in a subsequent round of interaction with *A. castellanii* for another six hours. This process was repeated five times and the resulting passaged strains were then named Pb18-Ac. The Pb18 strain, cultured in PYG for six hours at 28 °C, and then plated onto BHI-Sup was used as a control.

### *Galleria mellonella* infection

Wax moth larvae were raised in our lab and further details on the infection assay are described in the Supplemental Materials and Methods section. Shortly, groups of 12-16 individuals received an injection of 10 μl of PBS or yeast cell suspension (Pb18 or Pb18-Ac) at 10^6^ cells/mL in the hind left proleg. The groups of infected larvae were placed in Petri dishes, incubated at 37 °C and daily monitored for survival.

### Mouse infection and survival analysis

We infected isogenic 10-week-old BALB/c male mice with Pb18-Ac, using the non-passaged strain as negative control. The cells from each group were collected from BHI-sup plates after five days of culture, washed in PBS, counted, assessed for viability and diluted to the appropriated cell densities. The mice were anesthetized using a combination of 100 mg/kg of body weight ketamine and 10 mg/kg of body weight xylazine administered intraperitoneally. For infection, 10^6^ cells of either sample were intratracheally inoculated into two groups of 14 mice each. The animals were clinically monitored during 12 months after infection and moribund animals (defined by lethargy, dyspnea, and weight loss) were euthanized. The experiment was set up as a blind assay: the experimenters who infected and monitored the mice did not know which strain had been administered to each group until after the experiment finished. All mouse experiments were pre-approved by the Committee for Use of Animals in Research of the Catholic University of Brasília (protocol 017/14) in agreement with Brazilian laws for use of experimental animals and the Ethical Principles in Animal Research adopted by the Brazilian College for Control of Animal Experimentation.

### Quantitative RT-PCR of *P. brasiliensis* Pb18 and Pb18-Ac genes potentially involved in host-pathogen interaction

We analyzed the accumulation of selected transcripts previously involved in fungal response to macrophage or amoeba interaction by quantitative real time PCR as described in the Supplemental material and methods section.

### Flow cytometry analysis for the detection of alpha-glucan at the fungal cell wall

Pb18 and Pb18-Ac cells were paraformaldehyde-fixed and incubated with the anti-α glucan antibody MOPC 104E (Sigma) and then with a secondary anti-IgM conjugated with Alexa^®^ fluor 488. After washing, cell suspensions were analyzed in a BD LSR Fortessa flow cytometer. The resulting data were analyzed using FlowJo software.

### Statistical analysis

All statistical analyses were performed using GraphPad Prism 8.0 (GraphPad software). Percentage phagocytosis and percentage of amoeba viability (% Dead amoeba) were evaluated using Fisher’s exact test. Survival curves were analyzed using log rank and Wilcoxon tests. For CFU experiments we used one-way ANOVA with Tukey’s multiple comparison test or unpaired t test when comparing only two samples. Quantitative PCR analysis was performed with unpaired ttests.

## Results

### 1. Multiple groups of potential predators are present in the environment in which *P. brasiliensis* lives

Initial microscopical analysis of cultures obtained from soil samples positive for *Paracoccidioides* DNA as schematically represented in Figure 1A revealed the presence of multiple potential predators, including several amoeba morphotypes, ciliates and nematodes (Figure 1B-I). Although ciliates and nematodes are known to predate *C. neoformans* [11, 12], we chose to focus on ameboid predators. After using limiting dilution to obtain plates that seemed to contain only one type of amoeba, we made several attempts to establish axenic or monoxenic cultures. We were not successful, however, even in the presence of several antibiotics. We isolated DNA from the different isolates and performed PCR using primers specific to *Amoebozoa* and for *Acanthamoeba* spp identification. Sequencing and comparison against GenBank revealed that we had isolated members of *Allovahlkampfia spelaea, Vermamoeba vermiformis* (formerly *Hartmannella vermiformis*) and *Acanthamoeba* spp (Sequences were deposited under BioProject 506281).

**Figure 1.**
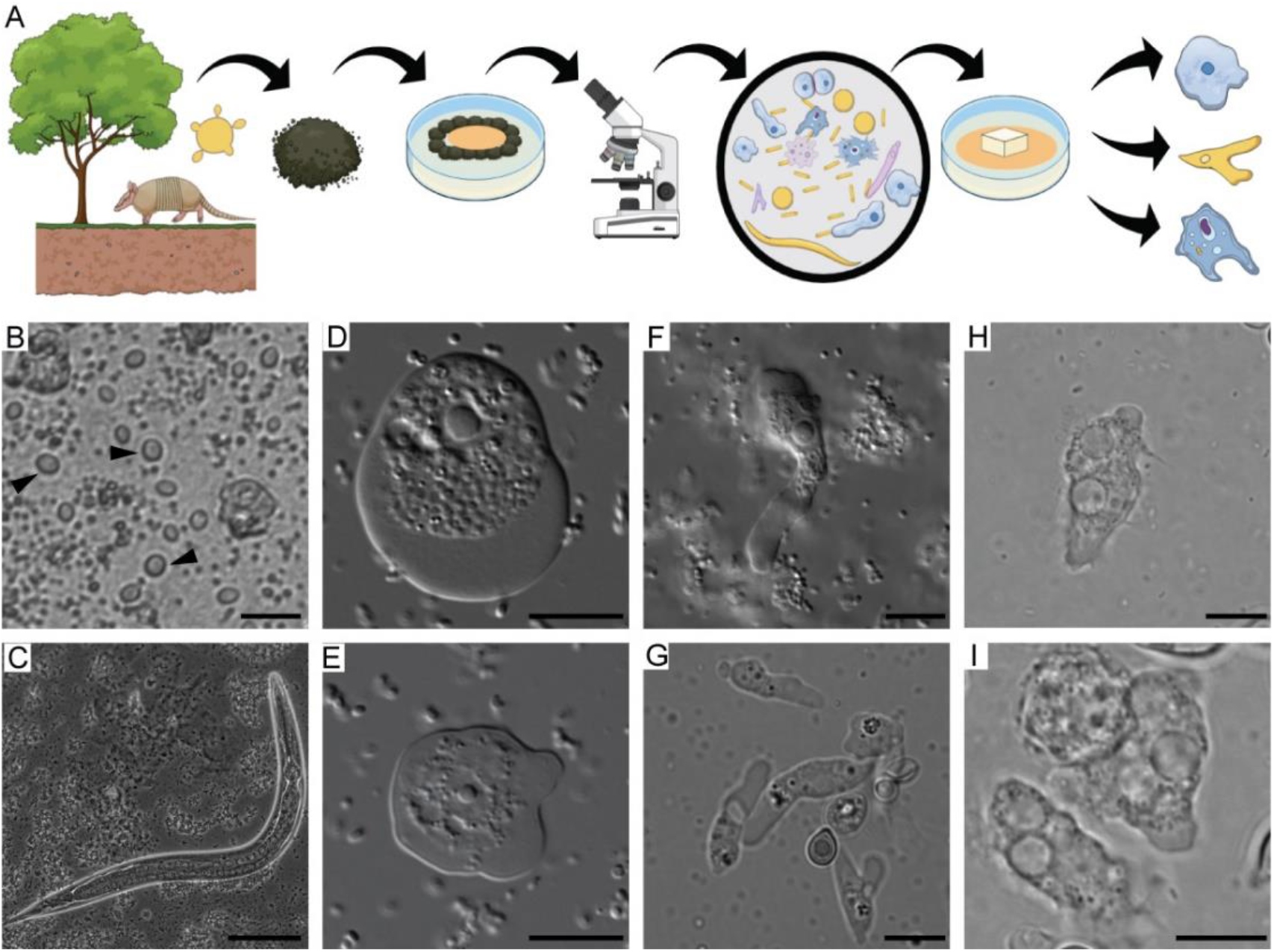
Soil organisms sharing the putative habitat of *P. brasiliensis*. A) Schematic representation of the soil amoeba isolation methodology. Soil samples from armadillo burrows positive for *P. brasiliensis* DNA were collected and used for the isolation of soil amoebae. The samples were plated in non-nutrient agar plates containing a bacterial lawn as food source and observed in an inverted microscope B) Bright field microscopy of trophozoites of ciliates (black arrow heads) present in a soil sample. Scale bar = 20 μm, C) Bright field microscopy of a nematode present in the soil sample. Scale bar = 50 μm, D) and E) DIC microscopy of trophozoites of *A. spelaea*. Scale bar = 10 μm. F and G) DIC microscopy of trophozoites of *V. vermiformis*. Scale bar = 10 μm. H and I) DIC microscopy of trophozoites of *Acanthamoeba* spp. Scale bar =10 μm.

### 2. Soil amoeba isolates interact with and kill *P. brasiliensis* yeast cells

We tested if these soil amoebae were able to phagocytose *P. brasiliensis* cells by co-incubating them for 24 h in PAS after adding antibiotic and removing most of the bacterial cells that were used to feed the amoebae. The three amoeba isolates were each able to phagocytose *P. brasiliensis* cells (Figure 2A), even in the presence of remaining bacterial cells from amoeba cultures, which are probably a preferential food source. We also observed that the isolated *Acanthamoeba* spp. had decreased viability after 24 h of co-incubation with *P. brasiliensis*, while the isolated *A. spelaea* was able to survive better in the presence of yeast cells (Figure 2B).

**Figure 2.**
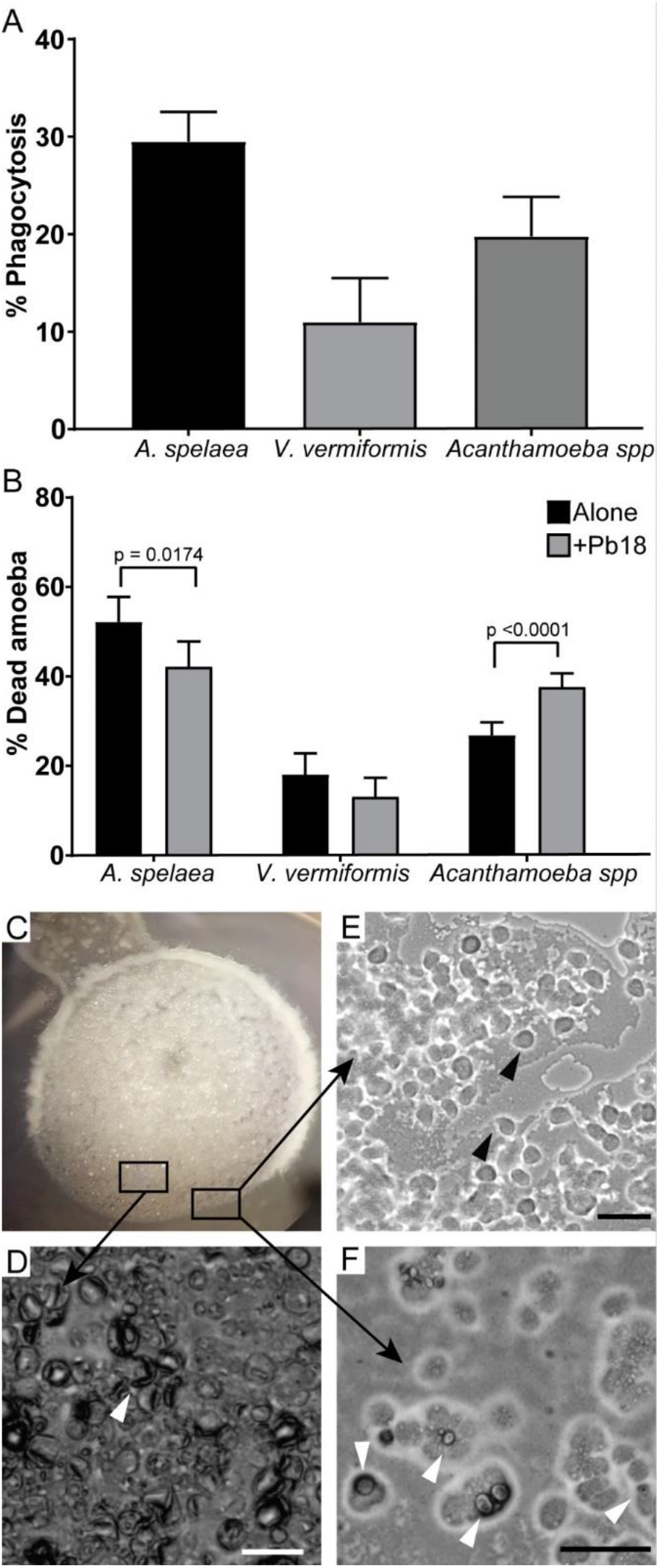
Interaction of *P. brasiliensis* Pb18 with amoebae isolated from soil of armadillo burrows positive for *P. brasiliensis*. The amoeba isolates were co-incubated with Pb18 previously dyed with pHrodo™ or FITC at a MOI of two at 25 °C for 24 h in liquid medium. A) Percentage of amoeba cells interacting with *P. brasiliensis* Pb18. After the interaction Pb18 cells were dyed using Uvitex 2B. B) Viability of the different amoeba isolates after 24 h of interaction with Pb18. A and D depicts the results of at least three independent experiments. At least 100 cells per replicate of each sample were counted for each assay. The bars represent 95% confidence intervals. C-F a suspension of *A. spelaea* cells was dropped next to a colony of *P. brasiliensis* cells in non-nutrient agar. The cells were co-incubated at 25 °C and examined daily in an inverted microscope. C) Macroscopic view of the fungal colony in a 35 mm plate. D) Microscopic view of the fungal cell lawn after seven days of interaction. E) Microscopic view of amoeba trophozoites growing in the periphery of the fungal lawn. F) Microscopic view of amoeba trophozoites interacting with a fungal cell. Scale bars = 50 μm. Black arrow heads indicate trophozoites. White arrow heads indicate fungal cells.

The presence of antibiotic resistant bacteria in the isolated amoebae cultures prevented us from evaluating *P. brasiliensis* viability after soil amoeba interaction by CFU counting. Due to the slow growth rate of this fungus, all the plates would be covered with bacteria before we could observe fungal colonies. To address this limitation, we performed predation assays in non-nutrient agar plate where *P. brasiliensis* lawns were confronted with soil amoeba isolates. We were able to observe a region of fungal cell clearance in the plates starting at 7 d of interaction with all the amoeba isolates as depicted in in Supplementary Figure 1. In Figure 2C-2F we can observe the interaction in solid medium of *A. spelaea* with *P. brasiliensis* cells. After seven days of co-incubation we could see many trophozoites mixed in the fungal cell lawn and around the colony (Figure 2D). Most fungal cells presented altered morphology resembling dead empty shells (Figure 2E) and we observed some fungal cells interacting with amoebae in Figure 2F. Additionally, after two weeks or more of interaction, we observed scarce fungal filamentation in the co-culture samples, possibly because most fungal cells were dead, while the control fungal spots without amoebae displayed intense filamentation.

We further evaluated the fungal interaction with three different species of amoeba in saline suspension after 24 h of interaction by TEM and SEM as presented in Figure 3. TEM analysis revealed internalized fungal cells and cell wall debris inside amoeba vacuoles (Figure 3A-J) and the presence of several extracellular empty fungal shells, some of them collapsed with little or no cytoplasm (Figure 3E-I). SEM confirmed the contact between the two microbes in all the interactions (Figure 3C, G, L). It should be noted that *V. vermiformis* cells can be considerably smaller than large *P. brasiliensis* mother cells (Figure 3K, L). Extreme morphological alterations were observed in most fungal cells upon interaction with the three amoebae. We observed many collapsed fungal cells, including mother cells with shrinking buds (Figure 3 G, H, L). Fungal cells presenting damage in their cell walls might explain the observation of empty cells walls in TEM (Figure 3 D, H, K). Altogether these results confirm the ability of amoebae to kill *Paracoccidioides* and strategies beyond fungal phagocytosis must be considered to explain how they do it.

**Figure 3.**
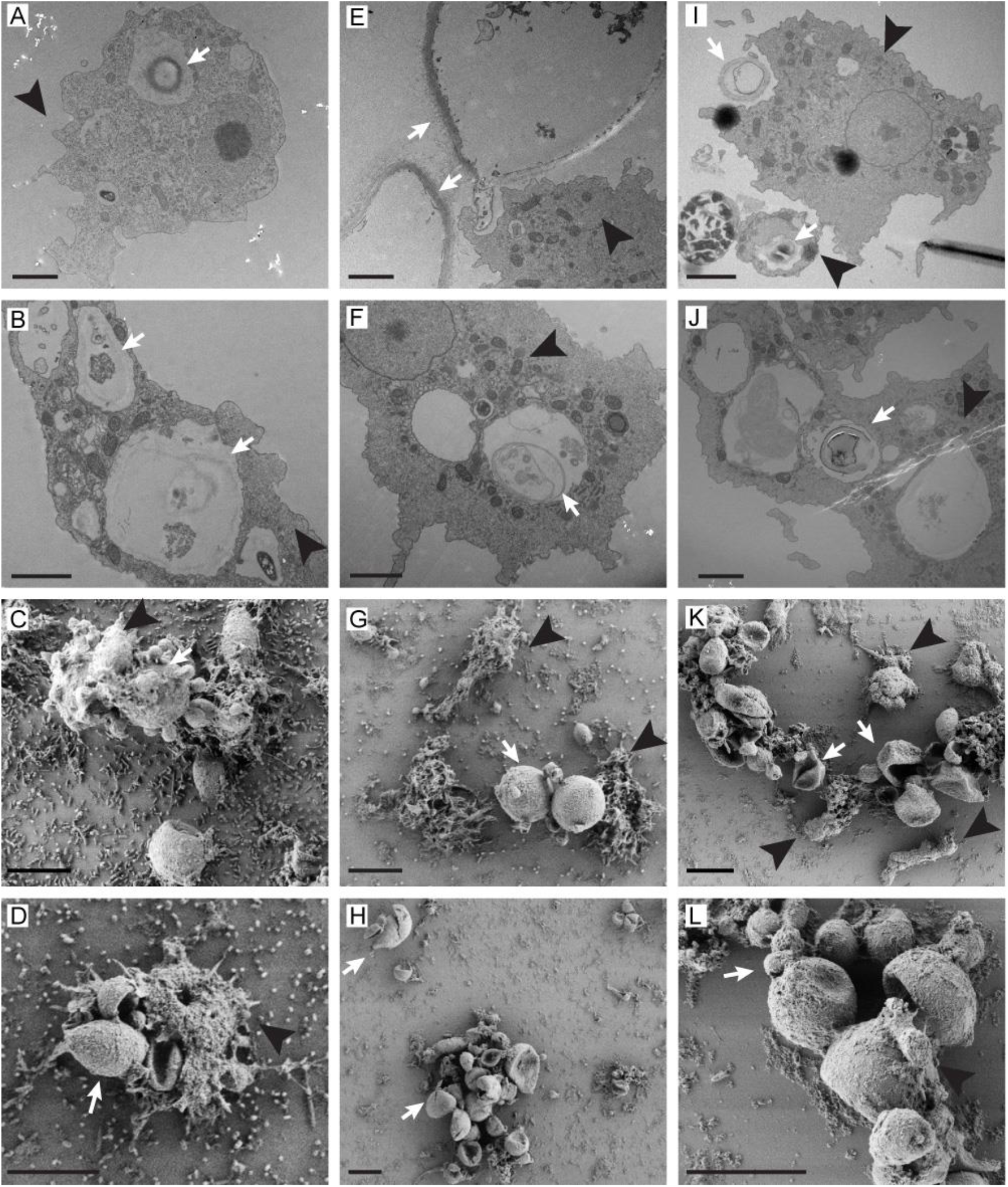
TEM and SEM analysis of the interaction of *P. brasiliensis* Pb18 cells and different soil amoebae. The isolates were co-incubated with Pb18 at a MOI of five at 25 °C for 24 h in PAS and fixed for microscopy. A-B) TEM of the interaction of *P. brasiliensis* with *A. spelaea*. Scale bars = 500 nm. C-D) SEM of the interaction of *P. brasiliensis* with *A. spelaea*. Scale bars = 10 μm. E-F) TEM of the interaction of *P. brasiliensis* with *Acanthamoeba spp*. Scale bars = 2 μm. G-H) SEM of the Interaction of *P. brasiliensis* with *Acanthamoeba spp*. Scale bars = 10 μm. I-J) TEM of the interaction of *P. brasiliensis* with *V. vermiformis*. Scale bars = 2 μm. K-L) SEM of the interaction of *P. brasiliensis* with *V. vermiformis*. Scale bars = 10 μm. White arrows indicate fungal cells, or their remains and black arrow heads indicate amoeba cells.

### 3. *Acanthamoeba castellanii* from axenic cultures can efficiently phagocytose and kill *P. brasiliensis* cells

Since the soil bacteria that remained in amoeba cultures was a third component of the microbial interaction system, and therefore a confounding factor, we decided to further evaluate the interaction of fungal cells with soil amoebae using axenized cultures of *A. castellanii*. Analysis of the co-culture of *A. castellanii* with Pb18 by light microscopy after Giemsa staining (Figure 4A-B), transmission electron microscopy (Figure 4C) and confocal microscopy (Figure 4D) revealed the interaction with and ingestion of yeast cells by amoebae. The cell wall-labelling dye Uvitex 2B (blue), which does not penetrate cells that are viable or not permeabilized, confirmed that some fungi were internalized. The black arrow in Figure 4D indicates an internalized yeast cell that is not labelled with CMFDA, which together with the irregular morphology suggests that this yeast cell is probably dead.

**Figure 4.**
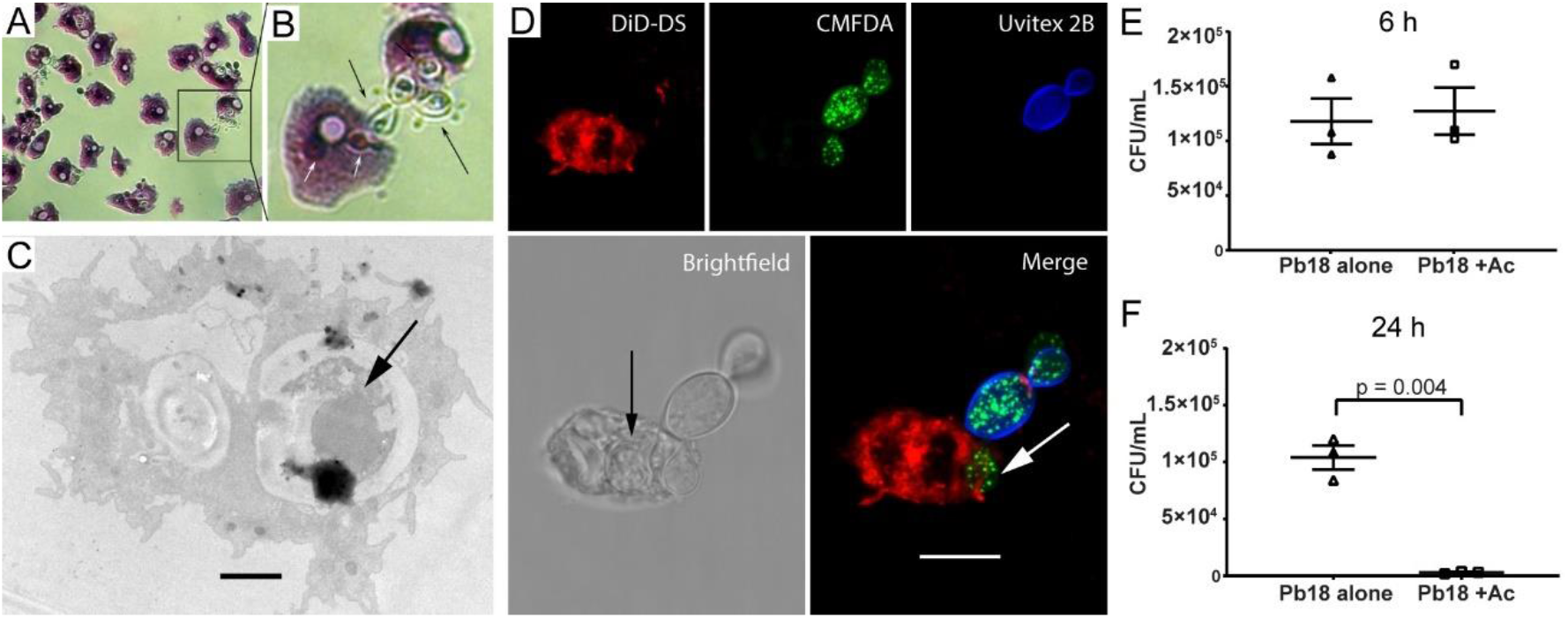
*P. brasiliensis* Pb18 interaction with an axenic *Acanthamoeba castellanii* strain. **A)** *P. brasiliensis* and *A. castellanii* were co-incubated at a MOI of one for one hour at 28 °C, and then stained with Giemsa and observed by light microscopy. **B)** Enlargement of the area depicted in the square region of panel A. **C)** TEM of the interaction of *A. castellanii* and Pb18 cells. Incubation was at a MOI of one for six hours at 28 °C and then fixed. The black arrow indicates an internalized *P. brasiliensis* (Scale bar = 2 μm). **D)** Confocal microscopy. *A. castellanii* was dyed with DiD-DS (red), while *P. brasiliensis* cells were labeled first with CMFDA (green), and after the interaction with Uvitex 2B (blue). The arrows show fungal cells inside an amoeba. (Scale bar = 10 μm). **E** and **F**) Survival of *P. brasiliensis* after interaction with *A. castellanii* interaction. Incubation was at a MOI of two at 28 °C for six (**E**) or 24 hours (**F**), using the fungus alone as a control. After the interaction amoeba cells were lysed and fungal cells were plated for CFU counting. The figure depicts the results of three independent experiments. The error bars represent standard error of the mean.

The percentage of phagocytosis of *P. brasiliensis* by *A. castellanii* was followed at different time intervals, from 30 minutes to 24 hours of interaction. It varied from 39% at 30 minutes to 68% at six hours (Supplementary Figure 2).

We also evaluated the outcome of amoeba predation by measuring fungal cell viability by CFU after six and 24 hours of interaction with *A. castellanii* cells. There was no significant reduction in fungal survival with or without amoeba at the earlier time point (Figure 4E), but the number of fungal CFUs was reduced by 90% after 24 hours of interaction (Figure 4F), indicating that *A. castellanii* was very efficient in fungal killing. On the other hand, the trypan blue exclusion assay on amoebae after interaction with *P. brasiliensis* showed that amoeba viability was barely affected by the fungus. We only found a small difference in their viability at the six-hour time point at 28 °C, but not at other times points at this temperature or at any time points at 37 °C when compared to the non-infected controls (Supplementary Figure 3A). To evaluate whether this effect resulted from a broader loss of virulence due to in vitro subculturing of the fungus, we tested the virulence of the same Pb18 isolate against J774 macrophages. In contrast with our observations with amoebae, we observed a significant decrease in macrophage viability after 24 (a 16% to 25% increase in dead cells) or 48 hours of interaction (22% to 33%) (Supplementary Figure 3A).

### 4. There are differences in the ability of different strains of *Paracoccidioides* spp. to survive interaction with amoebae

We also evaluated the interaction of *A. castellanii* with *P. lutzii* (Pb01) and *P. brasiliensis* T16B1, an isolate obtained from an armadillo. *A. castellanii* was able to internalize cells of the three different isolates of *Paracoccidioides* spp. at similar rates (Figure 5A). There was no difference in the ability of *P. lutzii* strain Pb01 relative to Pb18 to kill amoebae or to survive at six hours of interaction (Figure 5B-C). However, co-incubation with T16B1 resulted in a time dependent increase in the amoeba mortality in comparison to both other strains (Figure 5B and **Error! Reference source not found.**), while the other two isolates were able to induce a transient decrease in amoeba viability only at 6 hours of interaction. In addition, the armadillo isolate was also able to survive the interaction with amoebae better than the other two strains (Figure 5C). We observed an increase of roughly five-fold in the CFU of T16B1 after the interaction in comparison to the other two strains.

**Figure 5.**
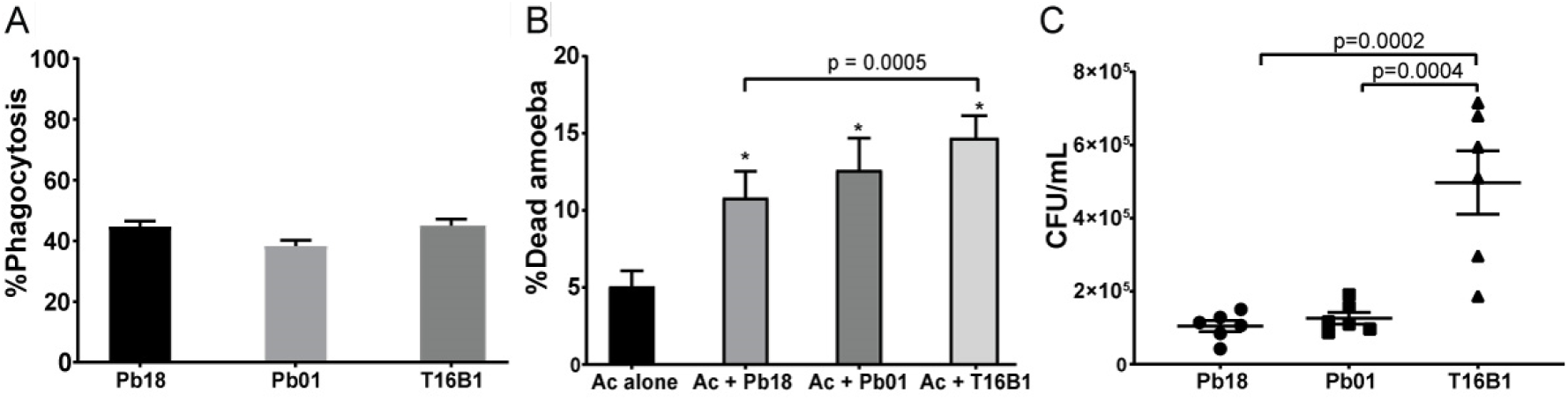
Interaction of *Paracoccidioides* spp strains with *A. castellanii* at six hours. Amoebae and three different strains of *Paracoccidiodes* spp (Pb18 – *P. brasiliensis*, Pb01 – *P. lutzii*, T16B1 – *P. brasiliensis* isolated from an armadillo spleen) were co-incubated at a MOI of two at 28 °C. **A)** Percentage of *A. castellanii* cells interacting with *Paracoccidioides* spp. The interaction was assessed by counting at least 100 phagocytes cells per replicate of each sample after Giemsa staining of the samples. The bars represent 95% confidence intervals. **B)** Viability of *A. castellanii* upon interaction with *Paracoccidioides* spp. The viability was assessed by counting at least 100 cells per replicate of each sample after staining with trypan blue. The bars represent means plus 95% confidence intervals. **C)** Survival of fungal cells from different strains of *Paracoccidioides* spp following interaction with amoebae. The error bars represent standard error of the mean. Figures depict the combined results of at least three independent experiments. *All the strains showed a significant difference in the % of dead amoebae at six hours relative to the control amoebae growing alone.

### 5. Sequential passages of interaction of *P. brasiliensis* with *A. castellanii* select for fungal cells with significant changes in their ability to survive and interact with different host models

We evaluated if sequential rounds of interaction of *P. brasiliensis* with amoebae were able to select fungal cells with increased virulence as previously reported for *H. capsulatum* [7]. We submitted the fungus to six hours of interaction with amoebae at 28 °C in PYG medium. The amoebae were then lysed and all interacting fungal cells (intracellular and extracellular) were collected and plated in solid BHI-sup medium. This procedure was repeated 5 additional times, resulting in Pb18-Ac strains. Both Pb18-Ac and Pb18 strains were used in co-cultures with *A. castellanii* and J774 macrophages and to infect *G. mellonella* and BALB/c mice. When comparing Pb18-Ac cells to the control strain, the phagocytosis decreased from 55.4% to 44.6% (Figure 6A), the proportion of dead amoebae increased from 10.8% to 15.9% (Figure 6B) and the number of fungal CFUs increased 2.5-fold (Figure 6C).

**Figure 6.**
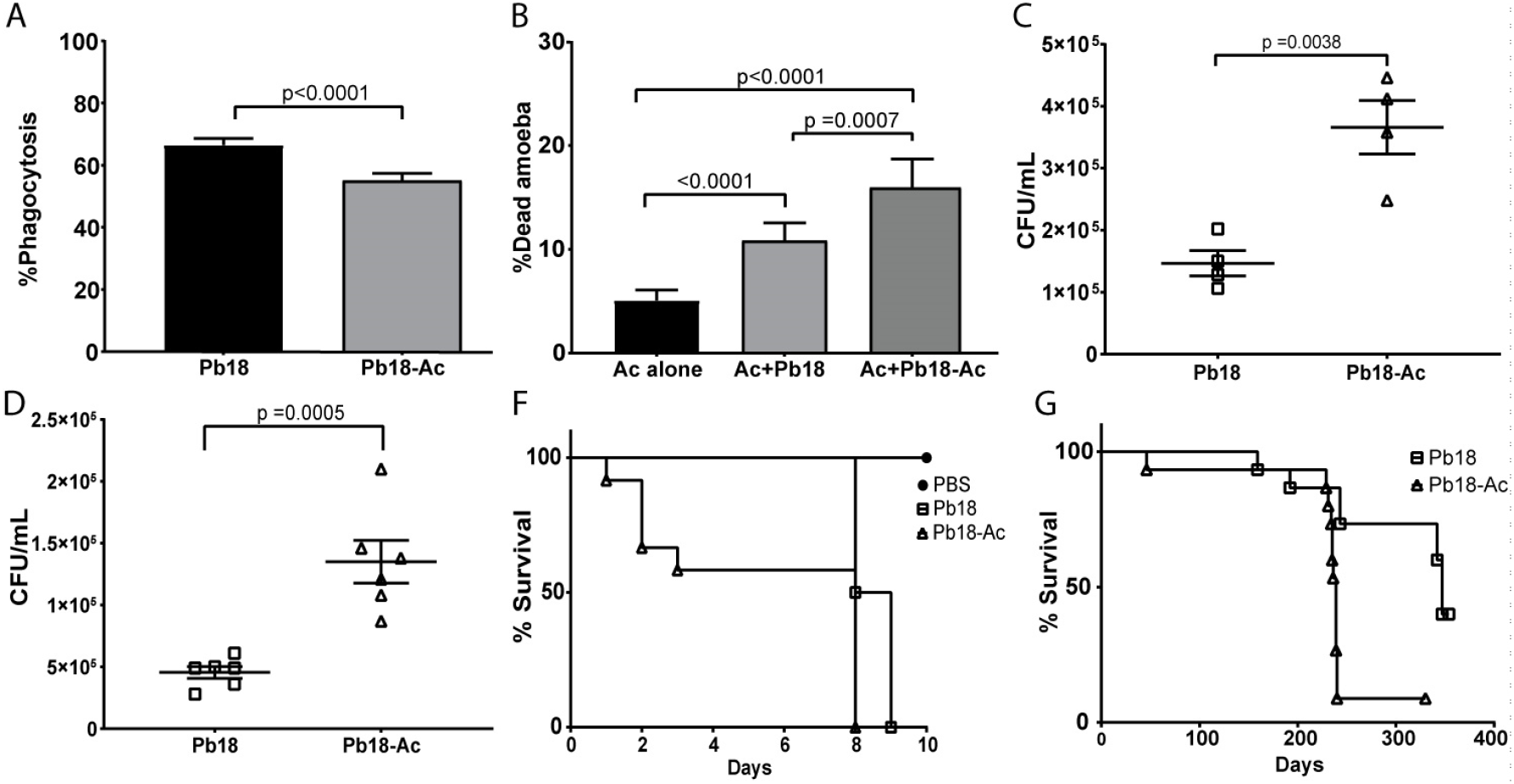
Effects of sequential passaging of Pb18 within amoebae, assessed in several models of infection. Pb18 and Pb18 Pb-Ac cells were co-incubated with *A. castellanii* at a MOI of two at 28°C for six hours. A) Percentage of *A. castellanii* cells interacting with Pb18 and Pb18-Ac. B) Viability of *A. castellanii* after six hours of interaction with Pb18 and Pb18-Ac. C) Survival of Pb18 and Pb18-Ac upon interaction with *A. castellanii*. D) Survival of Pb18 and Pb18 Pb-Ac upon interaction with J774 macrophages. E) Survival curve of *G. mellonella* infected with Pb18 and Pb18-Ac. The curve is representative of two biological replicates. P<0.0001 for the comparison of the survival curve of larvae infected with the two different strains (log-rank test). F) Survival curve of BALB/c mice infected with Pb18 or Pb18 Pb-Ac. Each group had 15 mice. p= 0.0003 for the comparison of the survival curve of mice infected with the two strains (log-rank test). A-D depict the combined results of at least two independent experiments. The bars represent means plus 95% confidence intervals in A and B and standard error mean in C and D.

Additionally, we also tested whether the changes in Pb-Ac interaction with amoebae could also be translated to other models. The number of recovered fungi in the wells with Pb18-Ac was significantly higher than the control strain after six hours of interaction with J774 macrophages (Figure 6D). Additionally, Pb18-Ac was also able to kill *G. mellonella* larvae and BALB/c mice significantly faster than the control strain (Figure 6E-F).

### 6. Sequential passaging of *P. brasiliensis* affects the accumulation of selected virulence transcripts and increases accumulation of cell wall a-glucans

Quantitative PCR analysis was carried out to search for changes in the levels of fungal transcripts that were previously shown to be modulated upon interaction with amoebae or macrophages [18–20]. No major changes were noticed in the accumulation of the transcripts of the selected genes between the two strains. The minor changes observed included a slight increase of HADH and HSP60 in Pb18-Ac (Figure 7B and E) and a slight decrease of *MS1* and *SOD1* expression (Figure 7C and D).

**Figure 7.**
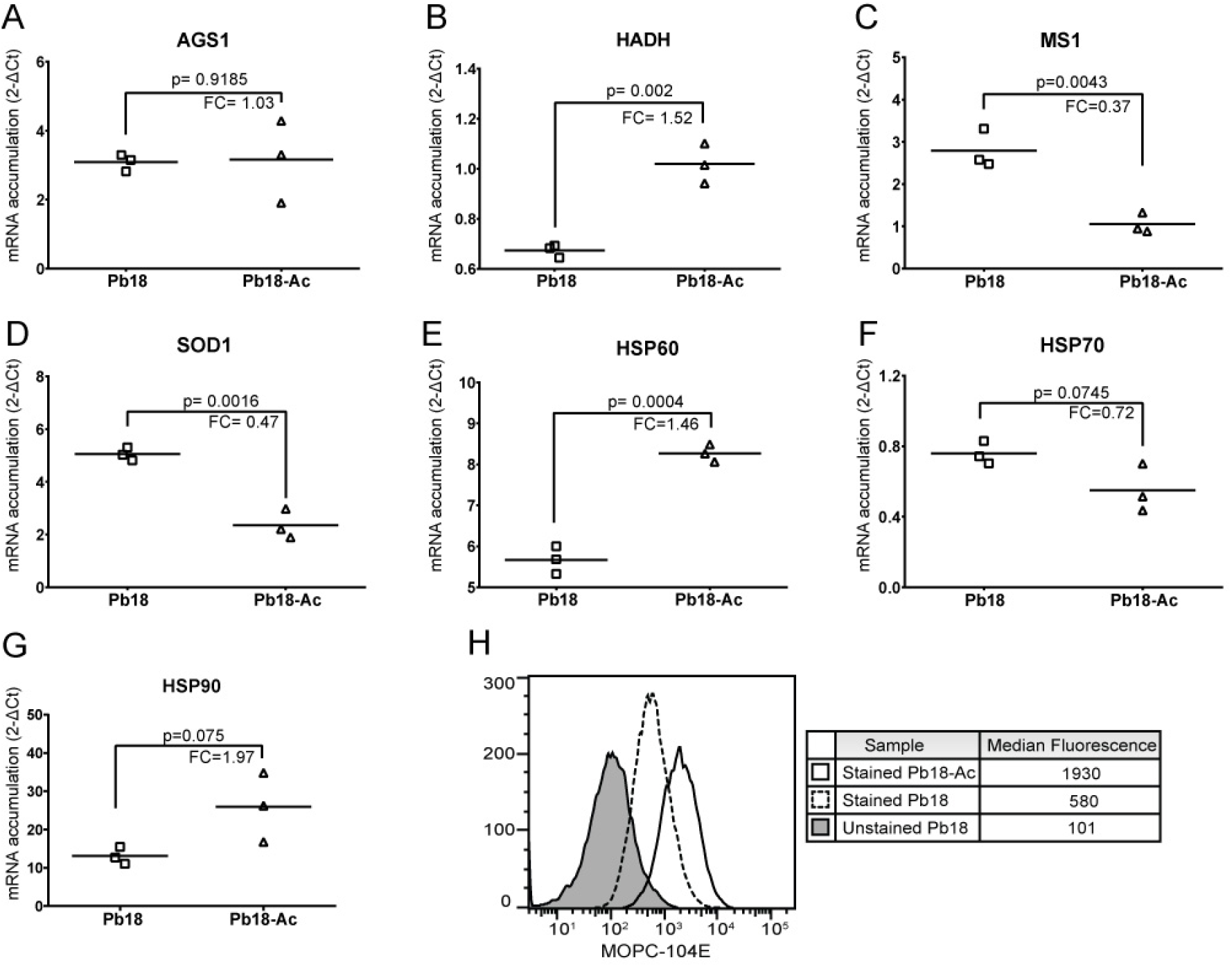
Modulation of Pb18 gene expression after passaging with amoebae. Transcript accumulation was determined by the comparative threshold method using the ΔCt value obtained after normalization with the constitutively expressed gene L34. Data are reported as individual 2^−ΔCt^ values of three independent experiments for each group and the bar represents their respective means. FC = *fold change in mRNA accumulation*, obtained as the ratio Pb18-Ac/Pb18. ** = p<0.01. ***= p<0.001. ns = p>0.05. **A)** AGS1: α-glucan synthase, **B)** HADH: Hydroxyacyl-CoA Dehydrogenase, **C)** MS1: malate synthase, **D)** SOD1: superoxide dismutase 1, **E)** HSP60: Heat shock protein 60, **F)** HSP70: Heat shock protein 70, **G)** HSP90: Heat shock protein 90. H) Cell surface staining of α glucan in the surface of Pb18 and Pb18-Ac.

As we observed a decrease in the percentage of phagocytosis of *A. castellanii* co-incubated with the passaged strain, we evaluated if there were any changes in the fungal surface that might affect its internalization. For that, paraformaldehyde-fixed Pb18 and Pb18-Ac cells were incubated with the monoclonal antibody MOPC-104E, which binds to fungal α-glucans [21] and analyzed by flow cytometry. Figure 7H shows a 3.3-fold increase in the signal for α-(1,3)-glucan in the passaged cells relative to the non-passaged strain. As an increase in the accumulation of α-glucan synthase transcripts was not detected by qPCR (Figure 7A), these results suggest that passaging through amoebae affect the content of α-(1,3)-glucans in *P. brasiliensis* cell wall through other mechanism that not mRNA accumulation.

## Discussion

*Paracoccidioides spp*. are thermodimorphic fungal pathogens that cause PCM, a systemic mycosis prevalent in Latin America [2]. Although this disease has been known for more than a century, there are still many unsolved questions about the ecology of its agents. Direct isolation of *Paracoccidiodes* spp. from soil is challenging and has been reported only a few times. However, detection of *Paracoccidioides* DNA from soil and aerosol samples is much more widely reported, suggesting that these fungi are saprophytes like those in the genera *Cryptococcus, Histoplasma* and *Blastomyces* [15, 22, 23].

In this study we analyzed the interaction of *Paracoccidioides* spp. with different soil amoebae. Our initial hypothesis was that the *Paracoccidioides* spp. virulence traits could have been selected by interactions with environmental predators such as amoebae and nematodes, as previously proposed for other soil-born fungal and bacterial pathogens. We performed interaction assays between *Paracoccidioides* spp cells and four amoebae, including three amoebae that we isolated from soil - *Acanthamoeba* spp, *A. spelaea, V. vermiformis* - and an axenic laboratory strain of *A. castellanii. Acanthamoeba* is a genus of soil amoeba that can cause keratitis and granulomatous amoebic encephalitis [24]. *A. spelaea* was first identified in 2009 and it was also involved in human keratitis, but there is little information about it in the literature [25]. *V. vermiformis* is frequently isolated from soil and water environments, including hospital tap water [26]. Interestingly, all three genera have previously been reported to harbor potentially pathogenic intracellular microbes such as *Legionella pneumophila* [27–29]. Additionally, there are several reports on the interaction of *A. castellanii* with different pathogenic fungi, and *V. vermiformis* has been shown to promote *Candida* spp. growth in tap water and conidial filamentation of *A. fumigatus* [7, 8, 10, 30–32]. In our experiments, all four amoebae were able to internalize and kill *Paracoccidioides cells* and non-axenic amoeba cultures were able to grow using fungal cells as their major food source. Different microscopy approaches have shown fungal internalization and many dead fungal cells or empty fungal cell walls after interaction with amoebae. Dead fungal cells presented severely altered morphology, many of which showing perforations in their surface. These results point to other mechanisms of *Paracoccidioides* spp killing by amoebae that do not depend solely on phagocytosis. *Paracoccidioides* spp. yeast cells are relatively large, and a mother cell with multiple buds is much larger than some of the amoebae we studied. Nevertheless, these amoebae efficiently killed the fungi. Our observations are supported by previous reports from the late 70’s of giant vampyrellid soil amoebae that perforated conidia and hyphae of several soil fungi such as *Cochliobolus sativus, Alternata alternaria* to feed upon their contents [33, 34]. This strategy bears a striking resemblance to vertebrate immune effector functions such as complement and granules from neutrophils, CD8+ T and NK cells. The selective pressure put on fungi by this amoeba feeding strategy could have led cell walls that are more resistant to the antimicrobial actions of these immune effectors.

At first, these results might contrast with what had been previously described for other pathogenic fungi such as *C. neoformans, Histoplasma capsulatum* and *Blastomyces dermatitidis* during the early 2000’s [7, 8]. However, our results are in accordance with previously published studies of fungal-amoeba interactions, in which *A. castellanii* was shown to efficiently use *C. neoformans* as a food source and was considered an important factor controlling fungi in the environment [35, 36]. More recently, Fu and Casadevall reported that divalent cations have the ability to increase amoeba survival and potentiate their fungal killing abilities [37]. Their work points to one of the factors that might influence the outcome of fungal-amoeba interaction assays. The studies published in the early 2000’s used PBS as the medium where interaction was assessed. In our assays we used PYG medium, which is supplemented with salts including CaCl2 and MgSO4. Additionally, the fungal cells used in our experiments were not submitted to regular passages through an animal host. Although *Paracoccidioides* spp is a primary pathogen, previous reports and our own experience have shown that prolonged in vitro subculture of these fungi leads to attenuation or loss of its virulence and that it can be restored by animal passaging [38, 39]. However, the same fungal strain was still able to kill a high proportion of macrophages in similar interaction conditions. These data suggest that despite the several similarities between amoebae and macrophages, there are important differences between these two cell host systems and/or that prolonged in vitro subculturing caused the fungal strain to lose virulence attributes that are more specific for its interaction with amoebae.

We further analyzed the interaction of *A. castellanii* with the sister species *P. lutzii* (Pb01) and with a *P. brasiliensis* strain isolated from an armadillo (T16B1). Regarding rates of internalization, and the ability to kill amoebae and to survive the interaction, Pb01 behaved identically to Pb18. However, the interaction of *A. castellanii* with T16B1 revealed that this strain was more efficient in surviving and killing the amoebae. Given that the armadillo strain was isolated about 7 years ago whereas Pb18 and Pb01 were isolated about 90 and 30 years ago, respectively [40, 41], these results point to the attenuation of *Paracoccidioides* spp. after prolonged in vitro subculturing. Our results are also compatible with previous work from other groups showing that *P. brasiliensis* armadillo isolates can be more virulent to mice and hamster models than some clinical strains submitted to prolonged in vitro culturing such as Pb18 [38, 42, 43].

Sequential passages of interaction with amoeba select for changes in the virulence of *P. brasiliensis* Pb18. When the control and passaged strains (Pb18-Ac) were used in new interaction assays with *A. castellanii*, Pb18-Ac cells were more efficient in evading phagocytosis, surviving the interaction and killing the amoebae. The decrease in the yeast internalization may be an important reason for increased survival of the fungus. As we did not separate internalized from non-internalized yeast for the fungal survival assays, the rate of internalization might have affected our measurement of the fungal survival. The passaged strain was also able to survive better the interaction with J774 macrophages and had an increased ability to kill *G. mellonella* larvae and mice, confirming that interaction with *A. castellanii* was able to select for broader changes in fugal virulence. These results are in accordance with what was described for *H. capsulatum* and *C. neoformans* upon their interaction with amoebae [7, 44]. Quantitative PCR of selected virulence genes between the two strains revealed downregulation of MS1 and SOD1, genes previously shown to be upregulated in response of *P. brasiliensis* to macrophage at six hours of interaction [18]. In contrast, HADH and HSP60 were upregulated. HADH is involved in beta-oxidation and in production of ergosterol precursors, and was previously shown to be upregulated in *P. brasiliensis* response to hypoxia [45]. This gene and others related to beta-oxidation were also shown to be upregulated in *C. neoformans* response to interaction with both amoeba and fungi [19]. Interestingly, despite no differences in the accumulation of α-glucan synthase transcript, we observed an increase in Pb18-Ac cell wall α-(1,3) glucans relative to Pb18. This change could explain the differences in fungal phagocytosis by amoeba. The outermost layer of α-(1,3) glucan in *H. capsulatum* cell wall has been shown to act as virulence factor by suppressing fungal recognition by host cells [21]. These results suggest similarities between fungal molecules that are recognized by phagocytosis receptors in amoebae and mammals. Overall, our results fit into the recently formulated amoeboid predator-fungal animal virulence hypothesis whereby there is a nexus of causation from selective pressure of amoeboid environmental predators and the evolution of fungal virulence against mammals [13]. We have shown that *Paracoccidioides* spp. may indeed interact with different amoebae species in its environment, and that soil protozoans, among other predators, could have a role as selective pressure for the emergence virulence traits in this genus, since amoebae can revert attenuation of its virulence from in vitro passaging. However, it should be noted that, although amoeba might indeed play an important role as a fungal predator in the soil and promote natural selection of virulence against animal host, there is many other potential predators and competitors in the soil. Investigation of fungal interactions with other soil inhabitants could shed light on many unsolved questions about the development of fungal pathogenesis.

## Supporting information

Supplemental Albuquerque et al. 2019

## Acknowledgments

This work was supported by grants from the Brazilian agencies, Conselho Nacional de Desenvolvimento Científico e Tecnológico (CNPq) and Fundacação de Apoio à Pesquisa do Distrito Federal (FAP-DF-Brazil). This study was also partially financed by scholarships from Coordenação de Aperfeiçoamento de Pessoal de Nível Superior (CAPES-Brazil, Finance code 001). We are grateful for the valuable help of several people along the development of this work including Barbara Smith, Thales D. Arantes, Raquel Theodoro, Marluce F. Hrycyk, Carlos Eduardo Winter, Jessica Ferrão, Gabriela Matos, Cristine Barreto, Izabella Monteiro Rizzi de Azevedo, Bianca Oliveira do Vale Lira, Calliandra de Souza and Jhones Dias.

## Conflict of Interest

The authors declare no conflicts of interest.

