## Supplemental Albuquerque et al. 2019 for "A hidden battle in the dirt: soil amoebae interactions with *Paracoccidioides* spp"

### Supplemental Figures

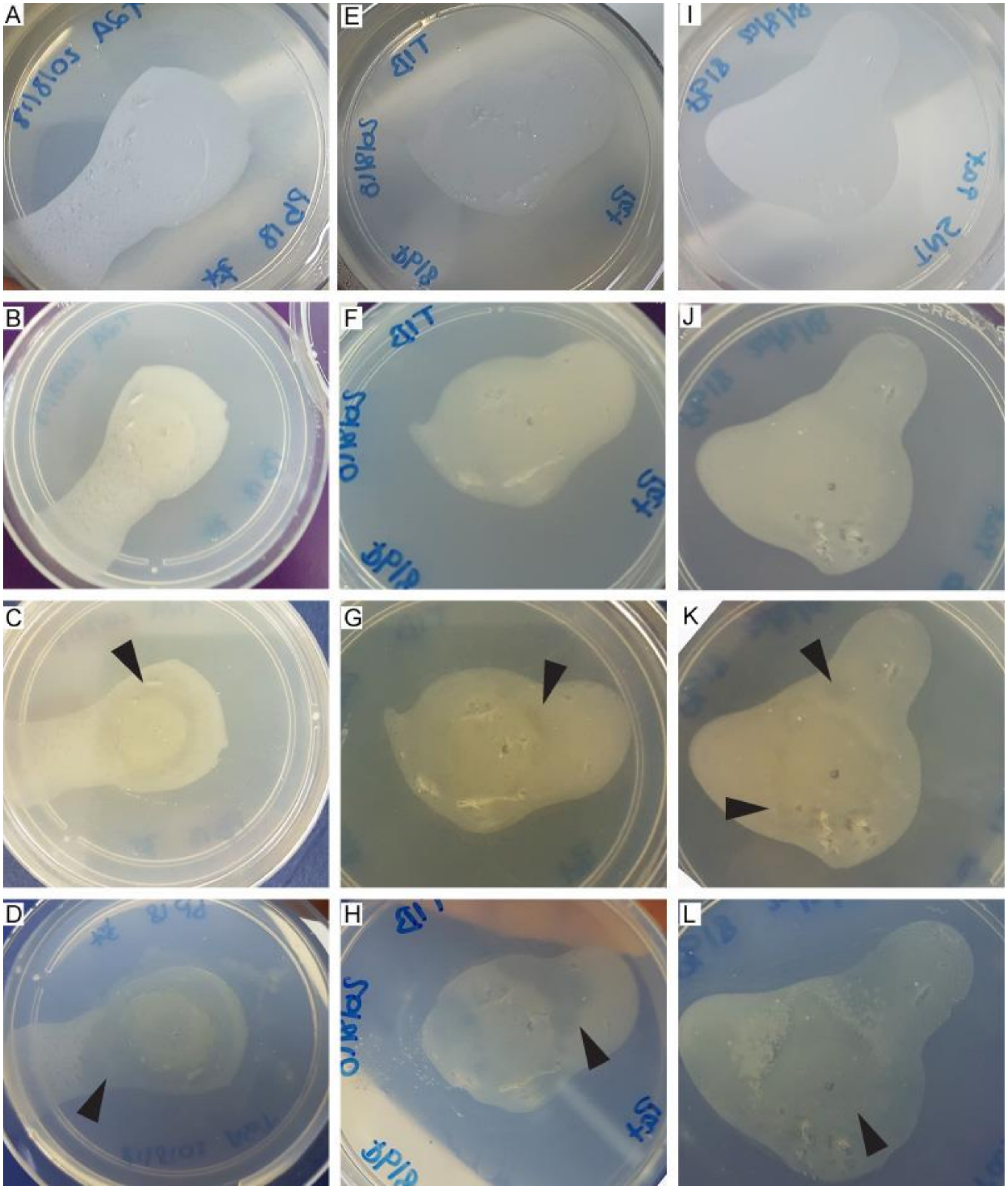

**Supplemental figure 1 - Interaction of *P. brasiliensis* Pb18 cells with soil amoeba isolates in solid plates of non-nutrient agar.**

A suspension of 1.5 $x10^7$ *P. brasiliensis* Pb18 cells was plated onto non-nutrient agar and spotted with $10^4$ cells of *Acanthamoeba* spp (panels A-D), A. *spelaea* (panels E-H) or *V. vermiformis* (panels I-L) in 10-microlitre aliquots directly in the middle of the fungal cell lawn. The plates were incubated at 25 °C for 19 days and inspected for the formation of lysis plates and fungal cell digestion at day 1 (panels A, E, I), day 3 (panels B, F, J), day 7 (panels C, G, K) and day 19 of interaction (panels D, H, L). Black arrowheads depict regions of fungal clearance.

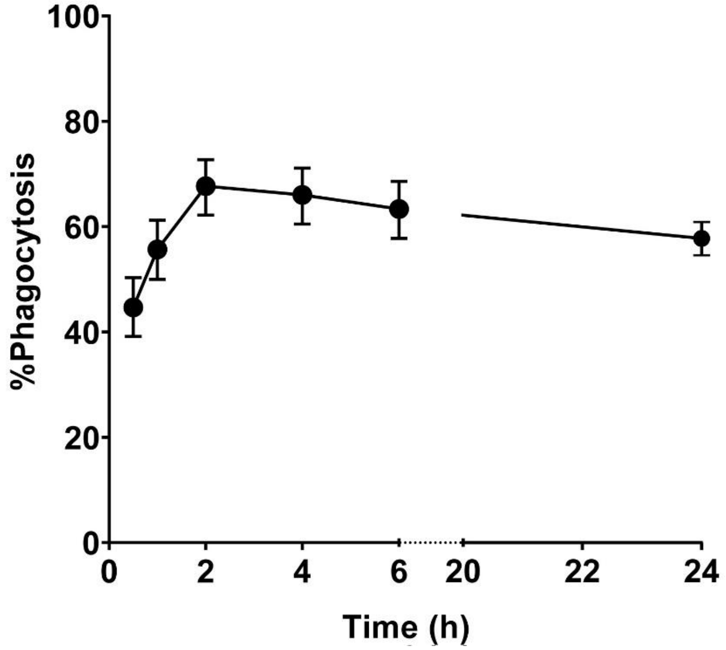


Supplemental figure 2 - Phagocytosis of *P. brasiliensis* P18 yeast cells by *A. castellanii*.

**Kinetics of phagocytosis of *P. brasiliensis* by *A. castellanii*.** Amoebae and *P. brasiliensis* yeast cells (CMFDA labeled) were co-incubated at a MOI of two. At each time point the percentage of phagocytosis was evaluated by fluorescence microscopy. A minimum of 300 amoebae per sample was analyzed to calculate the percentage of phagocytosis. The plot represents the results from three independent experiments each performed in triplicate. The error bars represent the 95% confidence interval.

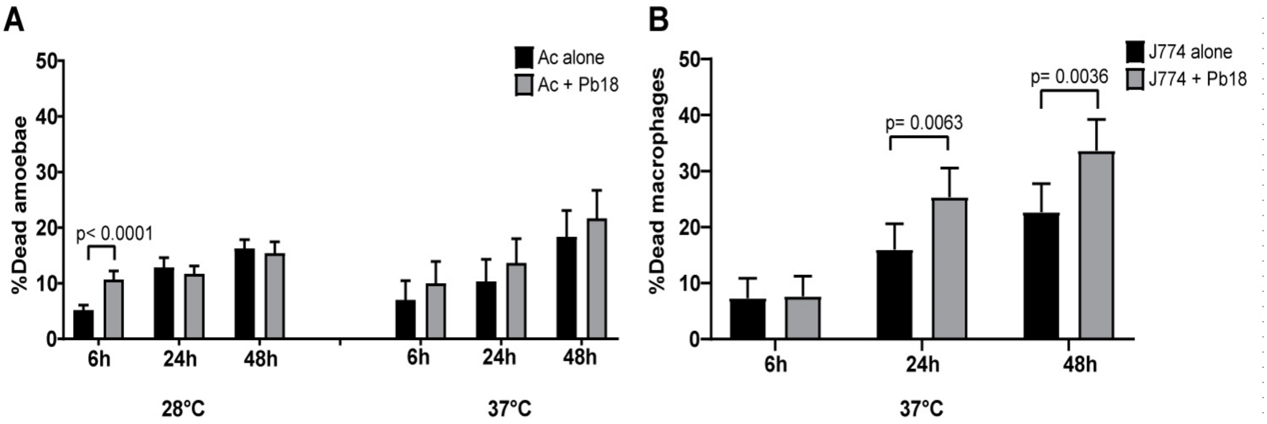


**Supplemental Figure 3 – Viability of *A. castellanii* and J774 macrophages after interaction with *P. brasiliensis* Pb18.**

A) Amoebae and yeast cells were co-incubated at a MOI of two at 28 °C or 37 °C for six, 24 and 48 hours. (B) J774 macrophages were incubated alone or in the presence of *P. brasiliensis* at 37 °C in a $CO_2$ incubator for six, 24 and 48h. Viability was assessed at each time point by counting at least 300 phagocytes cells per replicate after staining with trypan blue. The error bars indicate the 95% confidence interval.

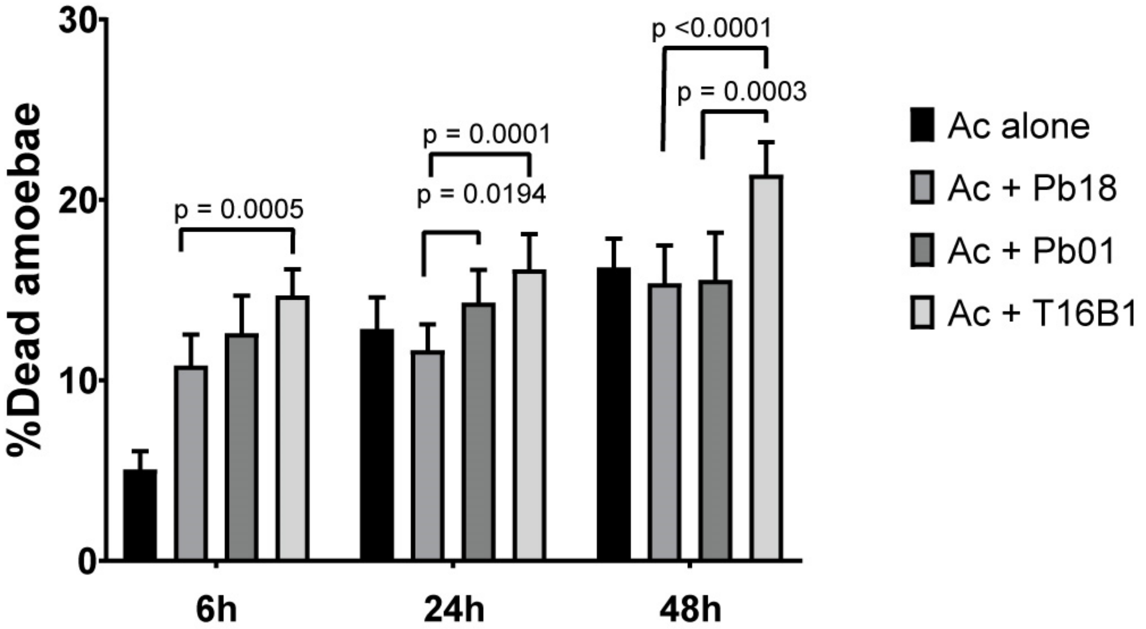


**Supplemental figure 4 –Viability of *A. castellanii* (Ac) after interaction with different *Paracoccidioides* spp strains.**

A) Amoebae were incubated alone or in the presence of *P. brasiliensis* Pb18, *P. lutzii* Pb01 or *P. brasiliensis* T16B1 yeast cells at a MOI of two at 28 °C or 37 °C for six, 24 and 48h. Viability was assessed at each time point by counting at least 300 phagocytes cells per replicate after staining with trypan blue. The error bars indicate the 95% confidence interval.

### Supplemental Materials and Methods

#### Fungal strains maintenance and preparation for interaction assays.

For our studies we used the *P. brasiliensis* clinical isolated isolate Pb18, *P. brasiliensis* isolate T16B1, isolated from the spleen of a nine-banded armadillo (*Dasypus novemcinctus*) (Arantes et al 2013) and the *P. lutzii* isolate Pb01. The yeast phase of these isolates was maintained by subculturing every seven days in Fava-Netto's medium medium (1.0 % w/v peptone, 0.5 % w/v yeast extract 0.3 % w/v proteose peptone, 0.5 % w/v beef extract, 0.5 % w/v NaCl, 4 % w/v glucose, and 1.4 % w/v agar, pH 7.2) or GPY medium (2% Glucose w/v, 1% Peptone w/v, 2% Agar w/v and 0.5% Yeast Extract) and incubating it at 37 °C. For experiments, five-day cultures were used. Before interaction assays, the fungal cells were collected, washed three times with PBS, and diluted to the appropriate cell densities. Only cultures with viability above 80%, measured with the viability dye, phloxine B (Sigma-Aldrich), were used.

#### DNA isolation for typing soil amoebae isolates

DNA extractions from soil amoebae were performed using the UNSET protocol (Hugo et al 1992) or QIAamp DNA Blood Mini Kit (Qiagen). Identification of amoeba isolates was performed by PCR using common amoeba primers AmeF977 (GATYAGATACCGTCGTAGTC) and AmeR1534 (TCTAAGRGCATCACAGACCTG) (Liang et al 2010) or *Acanthamoeba* specific primers JDP1 (GGCCCAGATCGTTTACCGTGAA)and JDP2 (TCTCACAAGCTGCTAGGGAGTCA) (Schroeder et al 2001). PCR fragments were purified and cloned into the TOPO™ TA vector (Thermo Fisher) and transformed into DH5-α *E. coli*. At least three plasmid clones were purified and Sanger-sequenced for each amoeba isolate. Sequences were blasted against GenBank and deposited under BioProject 506281.

*Paracoccidioides* spp. survival after interaction with amoebae

To assess survival of the fungus upon co-incubation with amoebae, the organisms were co-cultured for six or 24 h in 24-well plates, at a MOI of two. The cells were then detached from the plates and submitted to 5-8 passages through a 26-gauge syringe to lyse the amoebae. The remaining yeast cells were serially diluted and plated onto solid BHI supplemented with 4% horse serum, 5% conditioned medium of the Pb192 strain of *P. brasiliensis* (BHI-sup) (Castaneda et al 1988) and chloramphenicol (34 µg/mL). The plates were incubated at 37 °C for 7-10 days for colony counting. For each condition at least three wells were analyzed, and the experiments were performed at least three times on different days.

Confocal Microscopy

Sterile cover glasses were placed on six-well plates and 2 x $10^6$ amoeba cells in PYG medium were added to each well. After two hours of adhesion, amoebae were labelled with fluorescent dyes (DiD-DS for cell membrane or DDAO-SE for intracellular proteins). After that the amoebae were co-incubated with $10^7$ cells of *P. brasiliensis* (MOI of five) previously dyed with CMFDA. After two hours of interaction, the cover glasses were washed, fixed with cold methanol and mounted onto slides for confocal microscopy in a Leica SP5 microscope using a 63x NA 1.4 objective.

Transmission Electron Microscopy (TEM)

For electron microscopy, amoeba cells were co-incubated with *P. brasiliensis* yeast cells (MOI of five) for two or 24 h at 25 °C or 28 °C. Samples were fixed in 2.5% glutaraldehyde, 3 mM $MgCl_2$, 0.1 M sodium cacodylate buffer, pH 7.2 overnight at 4 °C. After rinsing with buffer, samples were postfixed in 1% osmium tetroxide in buffer (1 h) on ice in the dark followed by another rinse with 0.1 M sodium cacodylate buffer. Samples were left at 4 °C overnight in buffer, rinsed with 0.1 M maleate buffer, en bloc stained with 2% uranyl acetate (0.22 µm filtered, 1 h, in the dark) in 0.1 M maleate, dehydrated in a graded series of ethanol and propylene oxide, and embedded in Eponate 12 (Ted Pella) resin. Samples were polymerized at 60 °C overnight. Thin sections, 60 to 90 nm each, were cut with a diamond knife on the Reichert-Jung Ultracut E ultramicrotome and picked up with naked 200 mesh copper grids. Grids were stained with 2% uranyl acetate (aq.) followed by lead citrate and observed under a Philips CM120 TEM at 80 kV.Images were captured with an AMT XR80 high-resolution (16-bit) 8 Mpixel camera.

Scanning Electronic Microscopy (SEM)

Samples were fixed in 2.5% glutaraldehyde, 3 mM $MgCl_2$, in 0.1 M sodium cacodylate buffer, pH 7.2 overnight at 4 $^{o}$C. After rinsing with buffer, samples were postfixed in 1% osmium tetroxide in buffer (1 h) on ice in the dark followed by two distilled water rinses before dehydration in ethanol. Samples were dried for SEM with HMDS and mounted on carbon coated stubs, coated with 20 nm AuPd and imaged on a Leo FE-SEM at 1 kV.

Soil amoeba and *P. brasiliensis* predation assays

*P. brasiliensis* yeast cells were washed in PBS and big cell clumps were removed by passing the cells through 70-micron cell strainers or by multiple passages through 26-Gauge needles. After that, cell density was determined with a hemocytometer and a specific number of fungal cells were plated in the center of non-nutrient agar plates. After the fungal cell lawn dried, we aliquoted a suspension of the different soil amoeba isolates in the center of the plates. The plates were kept at 25 °C and examined daily for the presence of plaque-forming regions of fungal cell lysis.

*Galleria mellonella* infection

Wax moth larvae were kept in glass bottles in a dark environment in an incubator at 29 °C. The colony was maintained on an artificial diet consisting of portions of 500 g of Infant Cereal, 100 g of saccharose, 100 mL of glycerin, 100 g of honey and 100 mL of distilled water. Larvae weighing between 180 and 250 mg were used in the survival tests. Prior to each experiment larvae were collected, randomized into groups of 12-16 individuals and surface-cleaned with ethanol 70%. Each group received an injection of 10 µl of PBS or yeast cell suspension (Pb18 or Pb18-Ac) at $10^6$ cells/mL in the hind left proleg. All yeast suspensions contained ampicillin (20 mg/kg) to prevent infection with bacteria from the surface of the larva. The groups of infected larvae were placed in Petri dishes, incubated at 37 °C and daily monitored for survival.

#### Quantitative RT-PCR of *P. brasiliensis* Pb18 and Pb18-Ac genes potentially involved in host-pathogen interaction

Total RNA of Pb18 and Pb18-Ac cells was extracted using the Qiagen RNeasy Plant Minikit according to the supplied protocol, including treatment with the supplier's RNAse-free DNAse to avoid contamination with DNA. Samples were quantified in a Nanodrop spectrophotometer (Thermo Fisher). For cDNA synthesis, 2 µg of total RNA from each sample were reverse-transcribed using the High-Capacity cDNA Reverse Transcription Kit (Thermo Fisher) according to the manufacturer's instructions. The cDNAs were used as template for qPCR in triplicate using the Fast SYBR Green Master Mix (Thermo Fisher) with cycling conditions according to manufacturer's protocol, adapted for a reaction volume of 10 µL. The L34 transcript was used as endogenous control. The primers used had their amplification efficiency assessed by the standard curve method and are listed on Table S1. Changes in transcript abundance were quantified by the $2^{-\Delta Ct}$ method (Schmittgen & Livak, 2008), with fold-change determined as the ratio of values for each transcript between the non-passaged and passaged strains. The oligonucleotides used in these experiments are described in the following table.

| Gene | Forward Primer | Reverse Primer |
|---|---|---|
| Malate sintase (MS1) (PADG_04702) | TCAACTATCTCATGGAAGATGC | TCAACTATCTCATGGAAGATGC |
| 3-Hidroxi-acil-CoA desidrogenase (HADH) (PADG_01228) | GAGTTCGCCAACAAACTTCTCG | TGATCATGGAGCGGACTTGG |
| Alpha-glucan synthase (AGS1) (PADG_03169) | TCTGTGGCAACCTTGGGAGAC | TCCAGATTACTTGATGCTCAGTG |

| | | |
|---|---|---|
| Heat Shock Protein 60 (HSP60) (PADG_08369) | GATTACCAAGGACGGCGTTAC | TCTTGGAGGCAACGTCCTG |
| Heat Shock Protein 70 (HSP70) (PADG_00778) | TTCCTGGCTTGAAACACAGC | AACTCGCGGATTTTCGCTTC |
| Heat Shock Protein 90 (HSP90) (PADG_07715) | ATAAGACGCTGTCCAATGACTG | TTGGGCACGAAGAGGATGG |
| Superoxide dismutase 1 (SOD1) (PADG_07418) | AAGGCCGTCGCTGTTCTC | CTGTATGTGATGACGGTTGCG |
| 60S Ribosomal protein L34 (L34) (PADG_04402) | CTCCCGCGAATCCACAAC | ATGTGTTGGTGGGAGAGGAG |
